# Phase separation of ecDNA aggregates establishes *in-trans* contact domains boosting selective *MYC* regulatory interactions

**DOI:** 10.1101/2023.07.17.549291

**Authors:** Mattia Conte, Tommaso Matteuzzi, Andrea Esposito, Andrea M. Chiariello, Simona Bianco, Francesca Vercellone, Mario Nicodemi

## Abstract

Extrachromosomal DNAs (ecDNAs) are found in the nucleus of an array of human cancer cells where they can form clusters that were associated to oncogene overexpression, as they carry genes and *cis*-regulatory elements. Yet, the mechanisms of aggregation and gene amplification beyond copy-number effects remain mostly unclear. Here, we investigate, at the single molecule level, *MYC*-harboring ecDNAs of COLO320-DM colorectal cancer cells by use of a minimal polymer model of the interactions of ecDNA BRD4 binding sites and BRD4 molecules. We find that BRD4 induces ecDNAs phase separation, resulting in the self-assembly of clusters whose predicted structure is validated against HiChIP data (Hung et al., 2021). Clusters establish *in-trans* associated contact domains (I-TADs) enriched, beyond copy number, in regulatory contacts among specific ecDNA regions, encompassing its *PVT1-MYC* fusions but not its other canonical *MYC* copy. That explains why the fusions originate most of ecDNA *MYC* transcripts (Hung et al., 2021), and shows that ecDNA clustering *per se* is important but not sufficient to amplify oncogene expression beyond copy-number, reconciling opposite views on the role of clusters (Hung et al., 2021; Zhu et al., 2021; Purshouse et al. 2022). Regulatory contacts become strongly enriched as soon as half a dozen ecDNAs aggregate, then saturate because of steric hindrance, highlighting that even cells with few ecDNAs can experience pathogenic *MYC* upregulations. To help drug design and therapeutic applications, with the model we dissect the effects of JQ1, a BET inhibitor. We find that JQ1 reverses ecDNA phase separation hence abolishing I-TADs and extra regulatory contacts, explaining how in COLO320-DM cells it reduces *MYC* transcription (Hung et al., 2021).

## INTRODUCTION

Extrachromosomal DNAs (ecDNAs) are found in the nucleus of a broad range of human cancer cells where they can concur to oncogene amplification and correlate to poor prognosis and treatment efficacy (Kim et al., 2020; Nathanson et al., 2014, Turner et al., 2017). They are DNA rings, usually up to a few Mb in size and varying by tumour type, that can be composed of different oncogene harbouring chromosomal fragments (Gibaud et al., 2010; Shoshani et al., 2021; Rosswog et al., 2021; Verhaak et al., 2019; Vogt et al., 2004). Their mode of action, though, to organize and stimulate oncogene activation is still not fully understood despite their discovered role in cancer and their medical implications.

EcDNAs can be present in dozens or even hundreds of copies in the nucleus of cells, often with substantial heterogeneities across single cells (Chamorro González et al. 2023; Lange et al., 2022; Turner et al., 2017; Nathanson et al., 2014). They are marked by regions of accessible chromatin, by active histone signatures and include *cis*-regulatory elements, which could explain the high transcription levels of their genes, beyond copy number effects, also through local and chromosomal interactions (Morton et al. 2019; Wu et al., 2019). As ecDNAs are typically free to move in the nucleus, recent microscopy studies found, in an array of cancer cell types and primary tumours, that ecDNAs cluster with each other, also contacting chromosomal DNA, and colocalize with Pol-II (Adelman and Martin, 2021; Hung et al., 2021; Yi et al., 2021; Zhu et al., 2021). Those clusters include multiple rings that favor intermolecular enhancer-gene contacts and correlate with oncogene overexpression (Hung et al. 2021). Nevertheless, in some cases only specific copies of the oncogene on the ecDNA have a boost in transcription (Hung et al. 2021), hinting that ecDNA clustering *per se* is important but not sufficient to amplify expression. A different scenario was reported in other glioblastoma cell lines where it was observed that single rings are broadly dispersed in the nuclear volume and do not cluster (Purshouse et al. 2022). Additionally, it was found that the related increase of ecDNA oncogene transcription could be traced back to a change of copy number, hinting that no special mechanism is involved in ecDNAs activation and clustering, as after all the total number of their oncogenes, enhancers and mutual interactions is just proportional to the number of copies of ecDNAs (Purshouse et al. 2022). So, key questions remain open on how the clusters form and fold, how they enhance transcription beyond mere copy-number effects, and how they act selectively only at specific gene copies.

Here, we use polymer models to understand, at the single molecule level, the mechanisms whereby ecDNA clusters self-assemble and establish promiscuous contacts. As a case study we focus on a *MYC*-harboring ecDNA discovered in COLO320-DM colorectal cancer cells, whose molecular elements were comparatively well characterised (Hung et al., 2021). The ecDNA includes two copies of a *PVT1-MYC* fusion, an extra canonical copy of the *MYC* oncogene and an array of 47 distinct *cis*-regulatory elements associated with *MYC*. In COLO320-DM cells, the ecDNAs were shown to form clusters and to be tethered by the bromodomain and extraterminal domain (BET) protein BRD4, producing promiscuous contacts and resulting in a strong overexpression of the two *PVT1-MYC* fusions, but not of the other copy of *MYC* (Hung et al., 2021).

To investigate the link between ecDNA spatial organization and their functional role, we employ a minimal polymer model considering only the interactions of ecDNA BRD4 binding sites and BRD4 molecules. We find that BRD4 induces a phase separation of the rings, which form clusters while each ring undergoes a coil-globule transition. Next, we validate the model predicted three-dimensional (3D) structure of ecDNA clusters against available bulk H3K27ac HiChIP data (Hung et al., 2021), by remapping the model *in-cis* and *in-trans* contacts on the WT genome, hence disentangling the confounding effects deriving in the experiments from the overlapping genomic locations of the ecDNA genomic fragments.

We show that the ecDNA specific arrangement of BRD4 binding sites produces, via cluster phase separation, *in-trans* associated contact domains (I-TADs) enriched in promiscuous regulatory contacts among specific ecDNA regions. As the *PVT1-MYC* fusions are part of I-TADs, their interactions with regulatory elements become enriched beyond copy number effects. For steric hindrance among different rings, those effects plateau if the cluster size exceeds half a dozen rings. That shows, however, that even cells with comparatively few ecDNAs can be affected by strong oncogene amplifications. Because of the location of BRD4 binding sites, the other copy of *MYC* remains excluded from I-TADs and devoid of additional regulatory contacts. That explains the experimental finding that *PVT1-MYC* fusions are transcriptionally boosted by ecDNA clustering, whereas the other *MYC* is not.

Finally, we investigate the role of BET inhibitors, such as JQ1, that antagonize histone acetylated (H3K27ac) BRD4 chromatin binding sites for the bromodomain of BRD4 (Filippakopoulos et al. 2010; Nicodeme et al. 2010). In the model we find that above threshold concentrations of JQ1 reverse the ecDNAs structural transitions by turning off ecDNA-BRD4 interactions, hence abolishing the *in-trans* associated domains of the *PVT1-MYC* fusions and the related enhancement of regulatory contacts. That explains how JQ1 in COLO320-DM cells disrupts *MYC* transcription (Hung et al., 2021).

The picture emerging from our model reconciles the mentioned opposite scenarios on the very existence and mode of action of clusters of ecDNAs in distinct cancer cells by tracing back different observed behaviors to definite molecular features: in cells where ecDNAs do carry binding sites for specific chromatin organizing factors (e.g., in our model, BRD4 sites for BRD4 molecules) aggregation can occur by phase separation (Hung et al., 2021; Zhu et al., 2021), but in general clusters do not form (Purshouse et al. 2022).

Overall, our results clarify the mechanisms whereby ecDNA clusters boost selective oncogene activation, their pathogenic implications, and the potential role of BET inhibitors as anti-cancer drugs targeting BRD4-mediated oncogene-enhancer contacts. More generally, our findings shed light on the formation of nuclear aggregates of chromatin and other molecular elements, consistent with the experimental observations in the cell nucleus of condensates of factors, such as BRD4 and Mediator, and RNA polymerase II (Pol-II), and their involvement in gene activation (Barbieri et al. 2017; Cho et al., 2018; Chong et al., 2018; Conte et al. 2020; Sabari et al., 2018; Strom and Brangwynne, 2019).

## RESULTS

### Polymer model of the *MYC*-harboring ecDNA of COLO320-DM colorectal cancer cells

*MYC*-harboring ecDNAs of COLO320-DM colorectal cancer cells (Hung et al., 2021) are 4.3Mb long rings made of fragments from chromosome 6, 13, 16 and multiple fragments from chromosome 8 where the *PVT1* and *MYC* genes reside along with a broadly distributed array of BRD4 binding sites (**Fig. 1A**). Each ecDNA includes two copies of a *PVT1-MYC* fusion, a canonical *MYC* oncogene, an extra fragment of *PVT1*, and an array of *cis*-regulatory elements. In interphase nuclei, the ecDNAs form clusters, containing each 5 rings on average, tethered by BRD4 (Hung et al., 2021).

**Figure 1.**
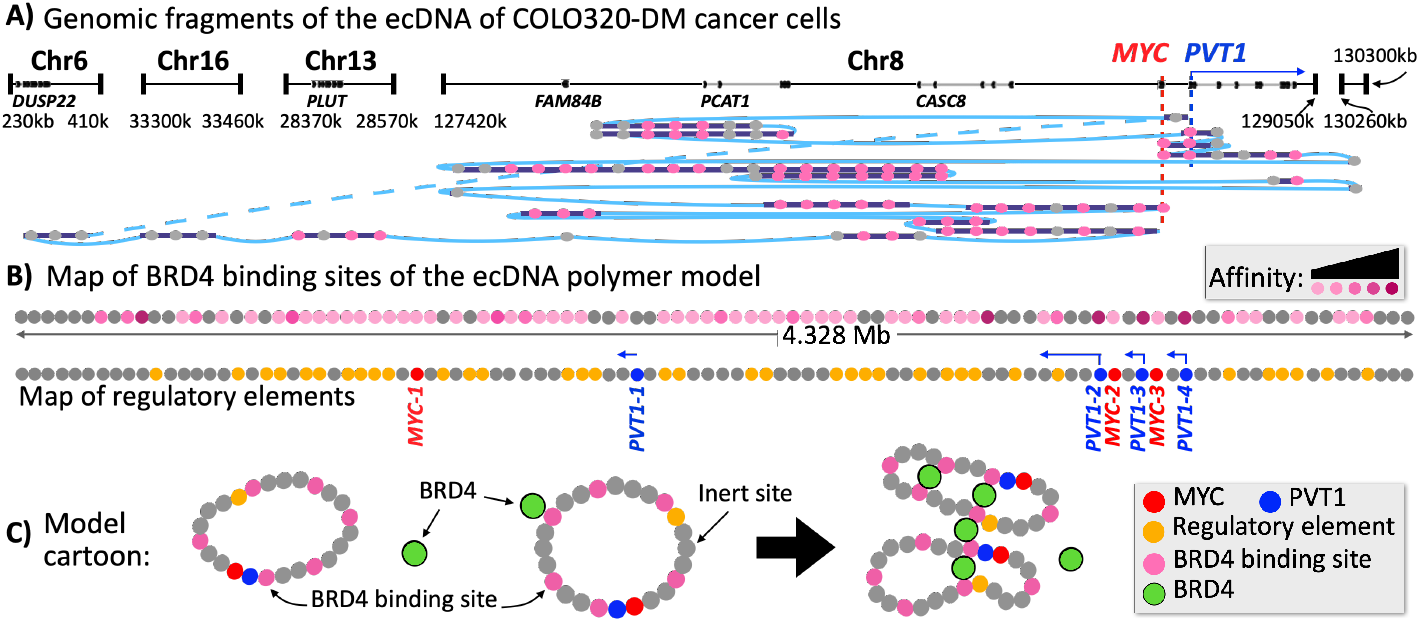
Ring polymer model of ecDNA of COLO320-DM colorectal cancer cells. **A)** The genomic arrangement of chromosomal fragments of the *MYC*-harboring ecDNA of COLO320-DM colorectal cancer cells and of its BRD4 binding sites (magenta circles) is mapped at 50kb resolution (Hung et al., 2021). **B)** Our *Strings and Binders* (SBS) polymer model of the ecDNA consists of a ring of beads with BRD4 binding sites (magenta) and BRD4 molecules (green) that can bridge them. Here a linear map is shown of BRD4 sites of the polymer model of the ecDNA, magenta shades represent different affinity strengths. Below are mapped the copies of *MYC* (*MYC-1, -2* and *-3*, red), *PVT1* (*PVT1-1, -2, -3*, and *-4*, blue) and *cis*-regulatory elements (yellow) (Hung et al., 2021). **C)** A cartoon of the polymer model and how it folds by interaction with BRD4 molecules.

To dissect the basic mechanisms of ecDNA aggregation and folding, we consider a polymer model that includes minimal molecular elements: the rings with their BRD4 binding sites and BRD4 molecules. We employ a model that has been extensively used to investigate chromatin 3D architecture, the *Strings and Binders* (SBS) model (Nicodemi&Prisco, 2009; Barbieri et al., 2012; Barbieri et al., 2017; Bianco et al., 2018; Conte et al., 2020; 2022). In the SBS model, an ecDNA ring is represented as a self-avoiding chain of beads, each associated to a coarse-grained 50kb window of DNA (**Fig. 1B**,**C**). The BRD4 sites on the chain (magenta in **Fig. 1B**,**C**) can be bound and bridged by diffusing BRD4 molecules. The affinities of the binding sites are chosen proportionally to the strength of their corresponding ChIP-seq signal (Hung et al., 2021) (**Methods**) and set in the weak biochemical energy range (from 2.25K_B_T to 6.25K_B_T, different shades of magenta in **Fig. 1B**). As BRD4 is known to form complexes, for example via its extraterminal (ET) domain, with other DNA factors (e.g., Mediator), in the model we represent those complexes (green circles in **Fig. 1C**) rather than single BRD4 proteins. Below we illustrate the case where they can interact with each other as experimentally reported (Sabari et al., 2018), with a reciprocal affinity again in the weak biochemical range (3.1K_B_T). Nevertheless, the case with no interactions between BRD4 molecules returns analogous results (**Methods, Supplementary Fig.s**) and, more generally, we checked the robustness of our findings by exploring variants of the model, with different values of DNA-binder and binder-binder affinities (up to 15K_B_T), coarse-graining levels, etc. (**Methods**). The system is subject to Brownian dynamics, investigated by extensive Molecular Dynamics (MD) computer simulations (up to 10^3^ runs made per set of parameters); the model units of measurements (i.e., length and time) are mapped into physical units (respectively, meters and seconds) by standard MD conversions (**Methods**). In a simulation, a given set of rings and BRD4 binders, initially randomly dispersed in a spherical volume (a proxy for the nucleus), approaches its steady state under the laws of physics (**Methods**). On each ring we also mapped the reported (Hung et al., 2021) copies of *MYC* (named *MYC-1, -2* and *-3*, in red from left to right in **Fig. 1B**) and *PVT1* (*PVT1-1, -2, -3*, and *-4*, blue in **Fig. 1B**,**C**) along with their *cis*-regulatory elements (yellow in **Fig. 1B**,**C**) to dissect the contacts they form by the action of BRD4.

### Phase separation of ecDNAs establishes selective, *in-trans* associated contact domains

To investigate the 3D structure of the ecDNAs model, for sake of clarity, we consider first the case with only a single ring. For the mentioned values of the model parameters, we find that upon increasing the concentration of BRD4 above a threshold value of around 4 nmol/l, the ring undergoes a conformational phase transition from a *coil* to a *globular* state (**Fig. 2A**) (de Gennes, 1979), signaled by a sharp drop in its average gyration radius, R_g_ (**Fig. 2B**). In the coil state, the average distance and contact matrix of the ring are featureless beyond genomic proximity effects (top **Fig. 2E, F**) because it folds in random conformations (top **Fig. 2G**) as entropy dominates the system free energy. In the globular state, conversely, specific contacts and loops appear especially between strong BRD4 binding sites (bottom **Fig. 2E, F**) because BRD4 mediated interactions thermodynamically prevail and fold the ring in compact, non-random lumps having an average diameter of around 380nm (bottom **Fig. 2G**). While phase transitions are robust thermodynamic processes, their exact threshold value depends on model parameters, e.g., binding energies, but we checked its order of magnitude is similar across the affinities we explored (**Methods**).

**Figure 2.**
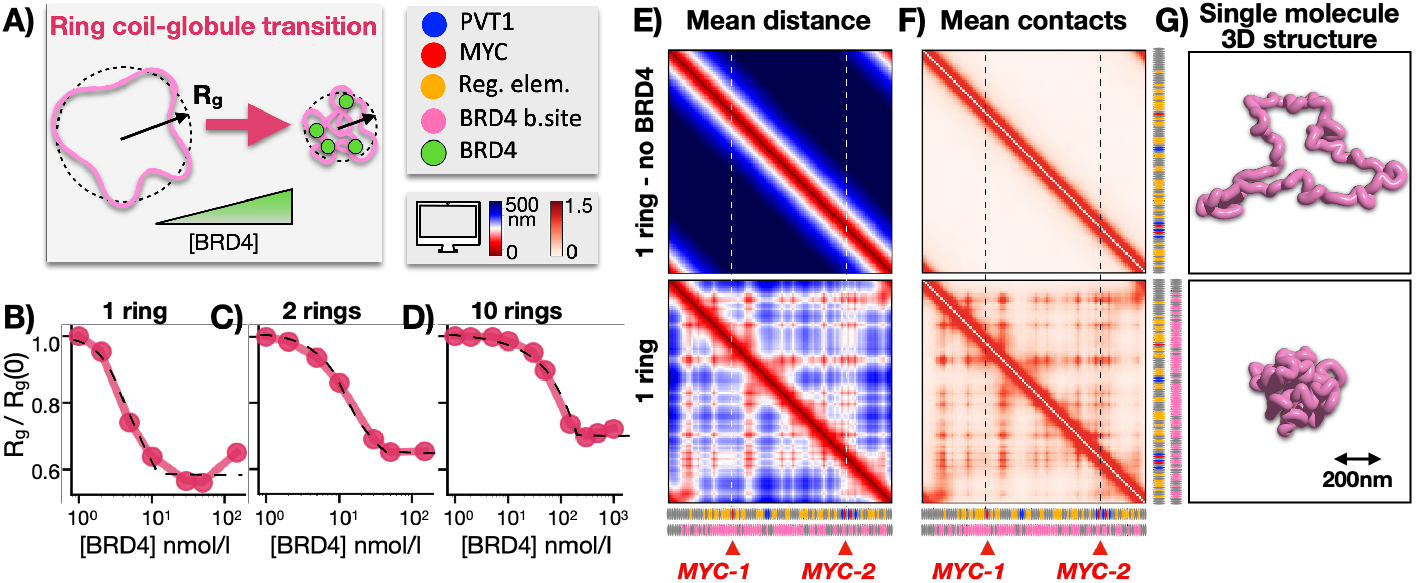
Single rings have a coil-globule transition driven by interactions with BRD4. **A)** In a model including only one ring, above a threshold concentration of BRD4, the polymer has a phase transition from a *coil* to a *globular state*. **B)** That is shown by a sharp drop of its average gyration radius. **C**,**D)** This structural change also occurs in a system with multiple rings. **E, F)** The average distance and contact matrices of a one ring model are shown in its coil (no BRD4, top) and globular state ([BRD4]=50nmol/l, bottom). **G)** A single-molecule ring 3D structure in the two phases.

In a system with multiple rings additional structural changes occur. We find that, as BRD4 concentration increases above a threshold, not only each polymer has a coil-globule transition (**Fig. 2C, D**), but the rings also *phase separate* into a single cluster by the action of BRD4 mediated interactions (**Fig. 3A**) (de Gennes, 1979), signaled by a sharp drop in the average steady-state distance of their centers of mass, D_CM_ (**Fig. 3B, C**). In the model, the average diameter of a cluster of 10 rings in a BRD4 concentration of 500nmol/l is around 850nm (**Fig. 3H**).

**Figure 3.**
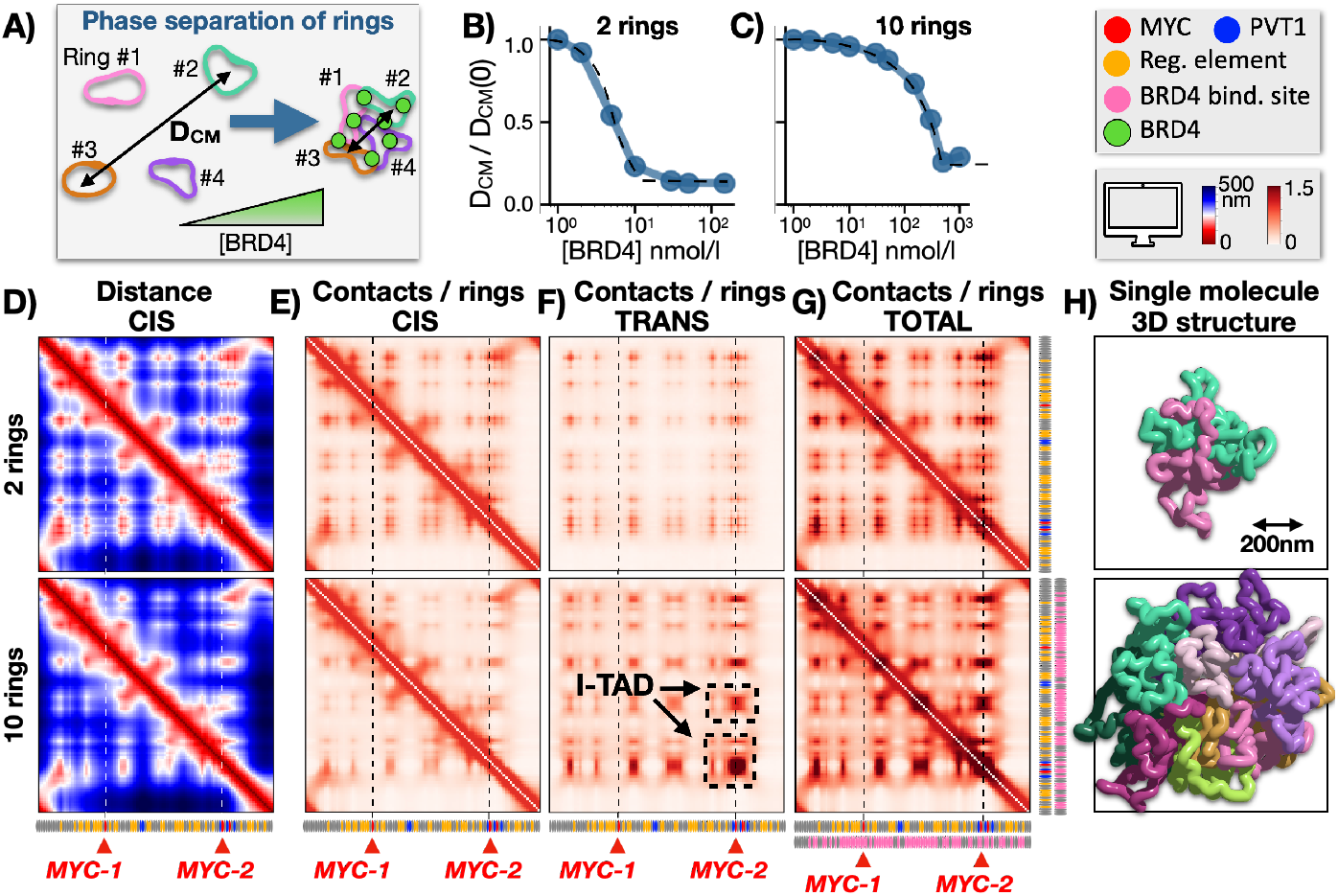
Phase separation of ecDNAs establishes *in-trans* associated contact domains. **A)** A system of multiple rings *phase separates* into a single cluster when BRD4 grows above a threshold concentration. **B**,**C)** That is marked by a sharp drop of the average distance of their centers of mass, in a system with resp. 2 and 10 rings. **D)** The average distance and **E)** the *in-cis*, **F)** *in-trans* and **G)** total contact matrices per polymer are shown in a system of 2 and 10 rings (resp. top and bottom) in the phase separated state. In a cluster of rings promiscuous contacts are not random, as specific *in-trans associated contact domains* (I-TADs, black dashed squares in the figure) appear between strong BRD4 binding sites. *MYC*-2/-3 and *PVT1*-2/-3/-4 are part of I-TADs and enriched of regulatory contacts, while *MYC*-1 and *PVT1*-1 are not, despite belonging to the cluster too. **H)** A single-molecule 3D structure of a cluster of 2 and 10 rings in the model phase separated state. The average diameter of a 10 rings cluster in 500nmol/l of BRD4 is around 850nm. The number of contacts plateaus in clusters larger than half a dozen rings because of steric hindrance.

We discover that, in the phase separated state, promiscuous contacts among rings are far from random, as specific interactions become established especially between regions enriched for strong BRD4 binding sites, as seen in the average contact matrix per ring (**Fig. 3F**, top and bottom panel respectively for 2 and 10 rings clusters) where specific *in-trans associated contact domains* (I-TADs) appear. I-TADs become more pronounced as the number of rings grows, up to saturate in clusters larger than half a dozen rings because of steric hindrance effects (**Fig. 3H**, see below). More in detail, as the number of rings increases, the contacts between a given pair of rings become on average weaker, yet total pairwise interactions *in-trans* per ring strongly grow since they can be formed combinatorially across many pairs (**Fig. 3G**). Interactions *in-cis* correspondingly decrease with respect to those of a single ring (**Fig. 3D, E**). Note that I-TADs result from a specific, non-uniform arrangement of BRD4 binding sites. For example, phase separation of homopolymers, which by definition have all equal beads, returns uniform contact matrices and no I-TADs, as shown in classical polymer studies (de Gennes 1979). In summary, promiscuous contacts are not uniformly distributed along an ecDNA, but are enriched in regions belonging to I-TADs, such as the two *PVT1-MYC* fusions, whereas the other copy of *MYC* and of *PVT1* (*MYC-1* and *PVT1-1*) remain excluded (**Fig. 3F, G**).

Finally, we characterize the nucleation kinetics of phase separation (de Gennes 1979; Brangwynne et al., 2015; Chiariello et al., 2016), i.e., how a single condensate emerges from an initially randomly dispersed set of polymers, by recording in time the average number of ring hubs and the average distance of the ring centers of mass. Those measures decay, after a characteristic time scale, τ_0_, respectively to 1 and to a plateau value, D_CM_ mentioned above (**Methods, Supplementary Fig**.). τ_0_ is an estimate of the self-assembly time of a single cluster: while its specific value depends on the system details, such as BRD4 affinities, we find it strongly grows with the number of rings in the system (**Methods**) (de Gennes 1979). For the mentioned model parameters, for example, we find that the condensation of a cluster of five rings takes around 8 hours, while ten rings take 22 hours (**Supplementary Fig**.). Hence, in a cell with a hundred ecDNAs the time scale of full aggregation can be so long that multiple, partial condensates are observed at shorter times. That rationalizes the finding that the average cluster in COLO320-DM cells, which include each on average 50 ecDNAs is made of around 5 rings (Hung et al. 2021). In the following, we focus again on the features of the final, single aggregate that is formed in the system steady-state.

Overall, our model shows that as soon as BRD4 concentration grows larger than a dozen nmol/l per ring, ecDNAs undergo structural transitions whereby they tend to fold on themselves (coil-globule transition) and cluster together by phase separation. The process spontaneously establishes specific *in-trans* associated contact domains, I-TADs, among the rings, determined by the genomic arrangements and strength of BRD4 binding sites. The resulting enrichment of contacts is highly site specific, depending on whether a given genomic region belongs to I-TADs (such as the two *PVT1-MYC* fusions) or it does not (such as *MYC-1* and *PVT1-1*).

### Model predicted 3D structure of ecDNA clusters is validated against HiChIP data

To test the model predicted 3D structure of ecDNA clusters against available experimental H3K27ac HiChIP bulk data (Hung et al., 2021), we remapped the rings’ genomic fragments (**Fig. 1A**) and their *in-cis* and *in-trans* interactions on the WT genome. While the model can disentangle the confounding effects deriving from the overlapping genomic arrangement of the ecDNA fragments, its WT remapped interactions can be directly compared to HiChIP (**Fig. 4A,B**): experimental and model contact matrices have overall similar patterns, with a Pearson and genomic-distance corrected Pearson correlation coefficients equal to r=0.7 and r’=0.6, which are comparatively high considering the minimal ingredients of the model.

**Figure 4.**
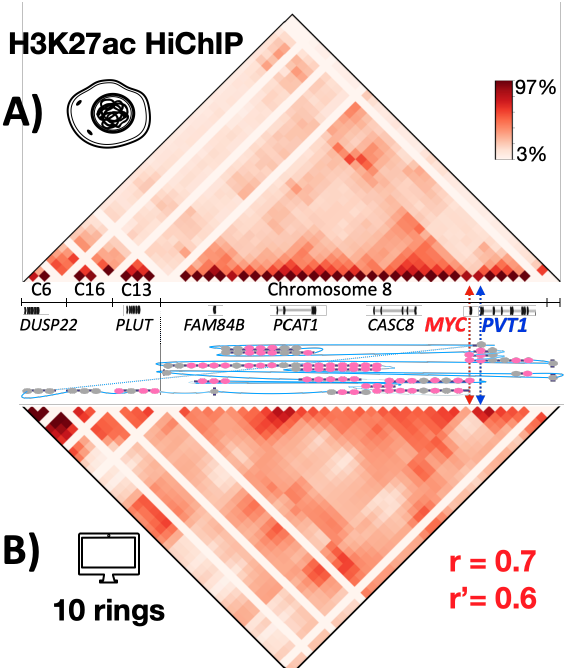
The model predicted 3D structure of clusters is validated by HiChIP data. **A)** H3K27ac HiChIP bulk data are shown for the ecDNA in COLO320-DM colorectal cancer cells (Hung et al., 2021). **B)** The total contact matrix per ring of phase separated clusters of 10 polymers (bottom Fig. 3G) is shown here remapped on the WT genome. The Pearson and genomic-distance corrected Pearson correlations of the two matrices are r=0.7 and r’=0.6, supporting the view that the model basic ingredients do capture important elements in the self-assembly of ecDNA clusters.

That supports the view that the basic ecDNA-BRD4 interaction mechanism, described by the model, does capture important elements in the self-assembly of ecDNA clusters and in the organization of their promiscuous contacts.

### ecDNA clusters boost promiscuous regulatory contacts of *PVT1-MYC* fusions, not of the canonical copy of *MYC*

To quantify the degree of interactions with regulatory elements established by the different genes in a cluster of ecDNAs, we computed virtual 4C contact profiles from the genes’ viewpoints (**Fig. 5**). As before, we consider the system in its phase separated state, report contacts normalized by the total number of rings and we take as a baseline the level of random contacts formed in absence of BRD4 (i.e., in the random coil state).

**Figure 5.**
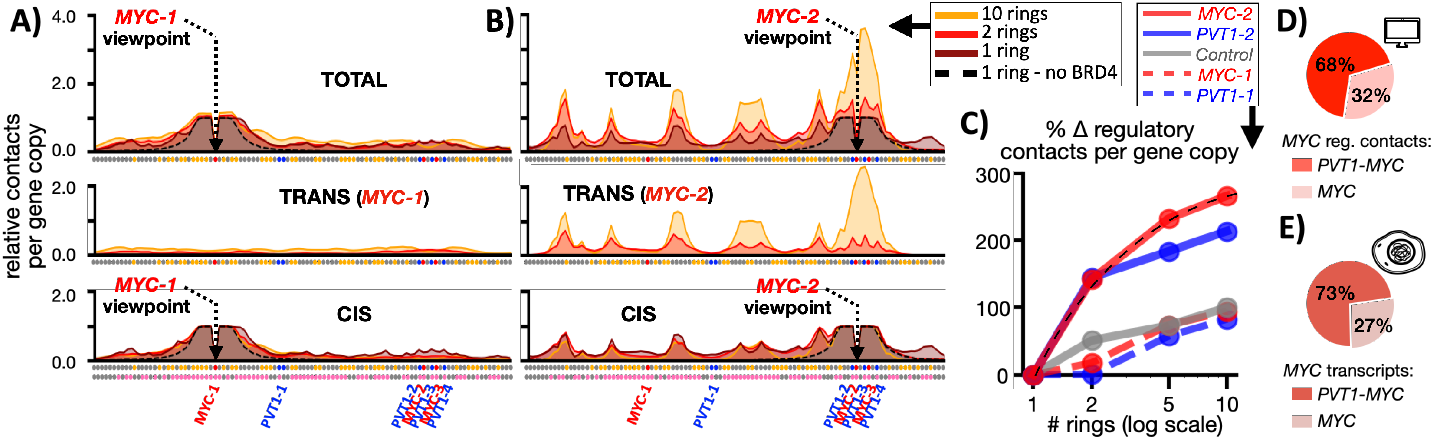
ecDNA clusters selectively boost *MYC*-2/-3 promiscuous regulatory contacts, not *MYC*-1. **A)** As the number of rings in a phase separated cluster increases, the virtual 4C contact profile (per ring) from the viewpoint of *MYC-1* only marginally grows above the baseline (black dashed line, contacts formed by *MYC-1* in a 1 ring model with no BRD4). **B)** Conversely, the promiscuous contacts of *MYC-2* markedly increase as it becomes part of I-TADs, because it is close to a strong BRD4 site (analogously *MYC-3*, not shown). That illustrates that cluster phase separation has different effects on the contacts of different gene copies on the ecDNA, depending on the underlying genomic arrangement and affinity of BRD4 sites. **C)** The change of regulatory contacts relative to regions devoid of BRD4 sites is shown as the number of rings in a cluster varies. *MYC-2* and *PVT1-2* have a threefold increase of contacts as the number of rings grows from 1 to 10. The change plateaus above half a dozen rings because of steric hindrance effects. *MYC-1* and *PVT1-1*, instead, are not different from the control, showing that ecDNA clusters selectively boosts beyond copy number effects *MYC-2* and *MYC-3*, but not *MYC-1* regulatory interactions. **D)** In a model system with 5 rings, the fraction of *MYC* regulatory contacts associated to *PVT1-MYC* fusions (i.e., *MYC-2* and *MYC-3*) is 68% of the total (including also two extra canonical copies of *MYC* to account for chromosomal genes). **E)** In COLO320-DM cells, where the mean number of ecDNAs in a cluster is five, *PVT1-MYC* fusions account for 73% of *MYC* transcripts (Hung et al. 2021). Those results combined hint that the oncogene transcription boost is controlled by the increase of its regulatory contacts.

We find that the total contacts formed by the canonical copy of *MYC, MYC-1*, has a modest increase over the baseline as the number of rings in a cluster grows (top **Fig. 5A**), despite their phase separation in a cluster, consistent with *MYC-1* exclusion from I-TADs (similarly for *PVT1-1*). The increase is driven by a slight increment of its interactions *in-trans* (middle **Fig. 5A**) while *in-cis* interactions weakly decline (bottom **Fig. 5A**). *MYC-2* has a radically different behavior: upon increasing the number of rings, its interactions *in-cis* slightly decline towards the baseline (bottom **Fig. 5B**), however correspondingly its contacts *in-trans* soar within its I-TADs (middle **Fig. 5B**), which is reflected by its total contacts (top **Fig. 5B**). That illustrates how phase separation of ecDNA has a major impact on the contacts of *MYC-2* well beyond copy-number effects (similarly for *MYC-3, PVT1-2, PVT1-3, PVT1-4*, **Supplementary Fig**.).

Next, we measured the variation of gene interactions (per gene copy) with regulatory sites mapped on the ecDNA, rather than total interactions. For a given gene we recorded the change of regulatory contacts relative to that of control beads devoid of BRD4 binding sites (gray beads on the polymer) when the number of rings in the system is increased (**Fig. 5C**). We find that while *MYC-1* and *PVT1-1* behave similarly to the control, *MYC-2* and *PVT1-2* (and analogously *MYC-3, PVT1-3* and *PVT1-4*) have a threefold surge in regulatory contacts as the number of ecDNAs in the cluster grows from 1 to 10. The change, though, tends to plateau above half a dozen rings because of the mentioned steric hindrance effects. In a model system with 5 rings, for example, we find (**Fig. 5D**) that the fraction of *MYC* regulatory contacts associated to the *PVT1-MYC* fusions is 68% of the total (including also two additional canonical copies of *MYC* to account for chromosomal genes). Interestingly, in COLO320-DM cells, where the mean number of ecDNAs in a cluster is five, the fraction of *MYC* transcripts associated to *PVT1-MYC* fusions is 73% of the total (Hung et al. 2021, **Fig. 5E**). Those findings hint that the oncogene transcriptional boost is controlled by a proportional increase of its regulatory contacts.

Taken together, our results explain why different copies of the *MYC* gene on an ecDNA undergo markedly different levels of upregulation upon the formation of ecDNA clusters. *MYC-2* and *MYC-3* are part of I-TADs produced by phase separation of the rings; hence their promiscuous regulatory interactions grow threefold larger than expected by copy numbers. Despite the formation of ecDNA clusters, *MYC-1* remains instead excluded from I-TADs, and its amplification derives merely from its multiple, independent copies in the cell. Such a picture rationalizes, for example, the experimental observation in COLO320-DM cells that three quarters of *MYC* transcripts originate from the *PVT1-MYC* fusions (Hung et al., 2021). Our model also points out that the aggregation of as little as half a dozen rings is sufficient to boost *MYC* upregulation, which then plateaus. Hence, oncogene amplifications can be significant even in cells carrying just a few ecDNAs, which could have substantial pathogenic implications.

### JQ1 reverses phase separation of ecDNAs and dissolves their contact domains

Finally, we investigate the impact of BET inhibitors (Filippakopoulos et al. 2010; Nicodeme et al. 2010) on the self-assembly of ecDNA clusters and on the resulting regulatory contacts. We focus on the effects of JQ1, a protein known to antagonize H3K27ac BRD4 chromatin binding sites for the bromodomain of BRD4. In our model, JQ1 is schematically represented by a new type of diffusing molecules that can bind BRD4 with a much higher affinity (14K_B_T) than binding sites along the rings. In the model, when a JQ1-BRD4 bond is formed, the resulting complex can no longer bind and bridge ecDNAs, to represent the unavailability of BRD4 bromodomains for DNA. Thus, JQ1 reduces the number of BRD4 molecules capable of bridging ecDNAs, and effectively turns off ecDNA-BRD4 interactions.

Specifically, for a given number of rings and a given initial concentration of BRD4, we investigated how the injection of JQ1 impacts the formation of contacts between *MYC* and regulatory elements. For sake of illustration, we focus on a system with 10 rings and [BRD4]=500nmol/l, and we vary the concentration of JQ1 (analogous results are found in a 5 rings system, **Supplementary Fig**.). We find that as JQ1 increases, *MYC-1* interactions (**Fig. 6A**) as well as the strong contact domains formed by *PVT1-MYC* fusions (**Fig. 6B**) go back to baseline levels. That is consistent with the experimentally reported effects of JQ1 in abolishing BRD4 chromatin occupancy (Loven et al., 2013). The effects of JQ1 are non-linear: the relative variation of gene interactions with regulatory sites (per gene copy) has a sigmoidal shape with its molar concentration (**Fig. 6C**). In fact, as soon as free BRD4 molecules are reduced below their critical threshold (**Fig. 3C**), phase separation is reversed, and ecDNA clusters and I-TADs dissolved. The inflection point of the sigmoid is where JQ1 has lowered free BRD4 at a value around such a threshold (**Fig. 6C**). In a model with five rings, for example, when the concentration of JQ1 goes from zero to 500nmole/l, we find (**Fig. 6D**) a 69% drop of *MYC* total regulatory contacts (including two additional canonical copies of *MYC* accounting for chromosomal genes). Correspondingly, in COLO320-DM cells when JQ1 grows from zero to 500nmole/l, the transcription probability of *MYC* was reported to fall by around 71% (Hung et al. 2021, **Fig. 5E**). That adds up to the results of the previous section in pointing out that *MYC* overexpression is mainly driven by regulatory contacts enhanced by ecDNA clusters beyond copy number.

**Figure 6.**
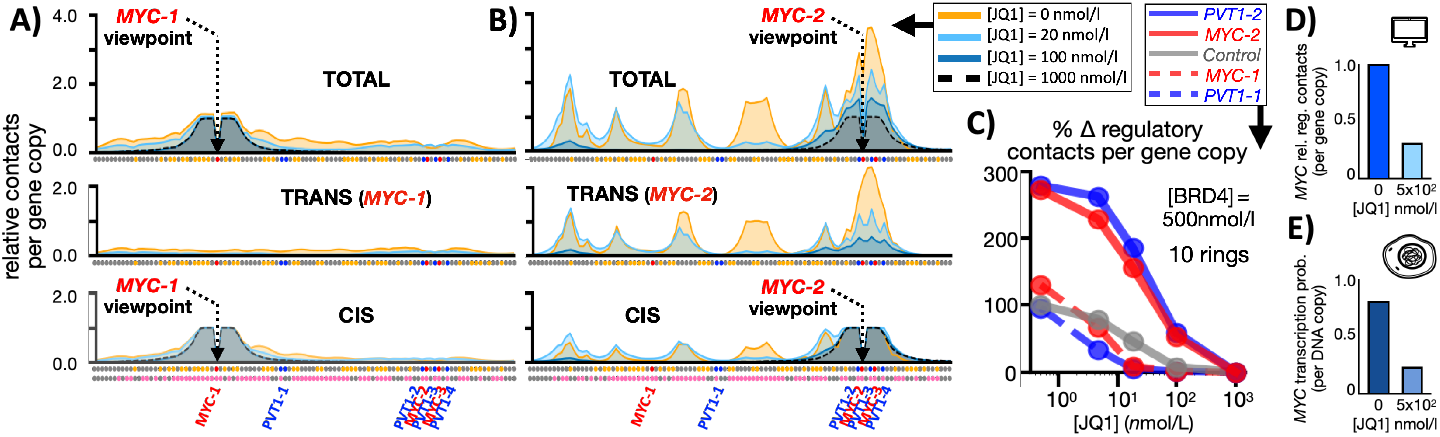
JQ1 reverses phase separation of ecDNAs and erases their contact domains. **A)** In our polymer model, the BET inhibitor JQ1 is represented as a new type of molecule that strongly antagonizes ecDNA BRD4 binding sites for BRD4, hence preventing the latter from bridging ecDNAs. The effects of JQ1 are illustrated in a system with 10 rings and [BRD4]=500nmol/l. As JQ1 concentration increases, the virtual 4C contact profile (per ring) from the *MYC-1* and, **B)**, *MYC-2* viewpoint collapse to the baseline. **C)** JQ1 has a non-linear impact on the relative variation of gene regulatory contacts (per gene copy): by reducing the concentration of free BRD4 below threshold it reverses phase separation, hence dissolving ecDNA clusters and their I-TADs. **D)** In a model with five rings, when the concentration of JQ1 goes from zero to 500nmole/l, *MYC* total regulatory contacts (including two additional canonical copies of *MYC* accounting for chromosomal genes) have a 69% drop. **E)** In COLO320-DM cells when JQ1 grows from zero to 500nmole/l, *MYC* transcription probability falls by around 71% (Hung et al. 2021). Our results combined hint that *MYC* overexpression, beyond copy number effects, is mainly driven by regulatory contacts enhanced by ecDNA clusters.

Overall, the model shows that JQ1 can reverse phase separation of ecDNA clusters by titrating out BRD4 molecules capable of bridging the rings. In turn, that erases *in-trans* associated domains and related enrichments in regulatory contacts. That explains the observation that high concentrations of the BRD4 inhibitor drastically reduce ecDNA *MYC* transcription (Hung et al., 2021). Because of molecular heterogeneities across single nuclei, however, different cells could have different responses to a given concentration of JQ1: cells carrying high enough levels of BRD4 would remain above threshold, thus maintaining *MYC* upregulation and its consequent pathogenic effects.

## DISCUSSION

To dissect the fundamental mechanisms of aggregation of *MYC*-harboring ecDNAs in COLO320-DM colorectal cancer cells, we introduced a minimal polymer model focusing on the interactions between BRD4 ecDNA binding sites and BRD4 molecules. Despite its simplicity, the model predicted 3D contact patterns and HiChIP data on ecDNA clusters (Hung et al., 2021) have good correlations, providing a validation of its basic molecular ingredients. The model also consistently explains a variety of experimental observations, ranging from the selective upregulation of the different copies of *MYC* to the role of JQ1 in disrupting *MYC* transcription.

The model shows that above threshold concentrations of BRD4 induce structural transitions in the rings. Their phase separation in clusters establishes *in-trans* associated contact domains, I-TADs, enriched in contacts among specific regions on the ecDNAs. As the two *PVT1-MYC* fusions are part of I-TADs, their interactions with ecDNA regulatory elements undergo a threefold increase, whereas the other gene copies *MYC-1* and *PVT1-1* do not, despite being part of the same aggregate. That rationalizes, for example, the experimental finding that three quarters of ecDNA *MYC* transcripts originate from *PVT1-MYC* fusions, rather than equally from all *MYC* copies (Hung et al., 2021). The model points out that the genomic arrangement and strength of BRD4 binding sites on an ecDNA is crucial to define the 3D structure of clusters: for example, the pattern of I-TADs is related to prominent BRD4 sites proximal to the promoter of *PVT1* on the ecDNA (Hung et al., 2021). Conversely, for example, a homopolymer would have a uniform contact matrix, i.e., no I-TADs (de Gennes 1979). The effects of I-TADs assembly become strong as soon as half a dozen ecDNA aggregates, showing that even cells with comparatively few rings could experience pathogenic *MYC* upregulations.

The model is, though, a simplified description of real ecDNAs, as other factors are likely to play a role, such as Pol-II and other chromatin organizers, which could contribute to establishing the 3D architecture (Pol-II and nascent *MYC* RNA, for example, are experimentally found to colocalize with ecDNA clusters (Hung et al., 2021)). In future works the model can be extended to incorporate those factors to help dissecting their roles, e.g., by monitoring how they would further increase correlations with HiChIP data. Nevertheless, the overall emerging scenario is robust as it is dictated by thermodynamics.

Additionally, our model clarifies that different species of ecDNAs in different cell types can have very distinct clustering behaviors according to their peculiar molecular nature: in cells where they carry binding sites for specific chromatin organizing factors (such as BRD4 sites for BRD4 molecules, as in our model) they can phase separate in clusters (Hung et al., 2021; Zhu et al., 2021), else in general they do not aggregate (Purshouse et al. 2022). That reconciles opposite experimental observations, in different cancer cells, about the very existence and functional importance of ecDNA clusters beyond copy-number effects.

Our approach can shed light on the potential of molecular factors to interfere at the molecular level with the BRD4 based clustering mechanism, beyond the discussed role of JQ1 in outcompeting ecDNAs for BET proteins. For example, one could study the impact on *MYC* regulatory contacts caused, e.g., (i) by factors interfering specifically with only the stronger BRD4 sites on the ecDNAs, rather than with BRD4 itself, hence preserving its natural functions, or (ii) by other genetic fragments carrying, for example, BRD4 sites but devoid of *MYC* and enhancer elements, as they would conglomerate into ecDNA clusters hence diluting overall *MYC* regulatory interactions. More generally, as our model can help testing and telling apart different scenarios on the mode of action of a variety of molecular factors, it could guide the design of novel anti-cancer drugs for targeted therapeutic applications.

## ACKNOWLEDGMENTS

MN acknowledges support from the National Institutes of Health Common Fund 4D Nucleome Program grant 5 1UM1HG011585-03, EU H2020 Marie Curie ITN n.813282, PNRR MUR M4C2 CN00000041 “National Center for Gene Therapy and Drugs based on RNA Technology” NextGenerationEU CUP E63C22000940007, MUR PRIN 2022 2022R8YXMR, and computer resources from INFN, CINECA, ENEA CRESCO/ ENEAGRID (Iannone et al., 2019) and Ibisco at the University of Naples.

## AUTHOR CONTRIBUTIONS

MN designed the project with help from MC. MN and MC developed the modelling with help from TM, AE, AMC, SB and FV. MC and TM ran computer simulations and performed data analyses with help from AE, AMC, SB and FV. MN and MC wrote the manuscript with inputs from all the authors.

## Notes

### Competing Interest Statement

The authors have declared no competing interest.

